# Functional reorganisation and recovery following cortical lesions: A study in macaque monkeys

**DOI:** 10.1101/202192

**Authors:** Matthew Ainsworth, Helen Browncross, Daniel J Mitchell, Anna S Mitchell, Richard E Passingham, Mark J Buckley, John Duncan, Andrew H Bell

## Abstract

Damage following traumatic brain injury or stroke can often extend beyond the boundaries of the initial insult and can lead to maladaptive cortical reorganisation. On the other hand, beneficial cortical reorganisation leading to recovery of function can also occur. Here, we used resting state FMRI (rsFMRI) to examine how functional connectivity in the macaque brain changed across time in response to lesions to the prefrontal cortex, and how this reorganisation correlated with changes in behaviour. Two monkeys were trained to perform location-based and object-based delayed match-to-sample tasks. We also collected rsFMRI data under general anaesthesia at two pre-lesion time-points, separated by 3-4 weeks. After two cycles of testing and scanning, the animals received a principal sulcus lesion followed by an additional 4 cycles of testing and scanning. Later, the same animals received a second lesion to the opposite hemisphere and additional cycles of testing and scanning.

Both animals showed a marked behavioural impairment following the first lesion, which was associated with a decrease in functional connectivity, predominantly within frontal-frontal networks in both hemispheres. Approximately 8 weeks following the lesion, performance improved, as did functional connectivity within these networks. Following the second lesion, functional connectivity again decreased and this was associated with a marginal behavioural deficit that did not recover.

Our data show that behavioural impairments reflect not just the removal of the lesioned area, but also disturbance to an extensive cortical network. This network can recover by restoring and/or strengthening pre-existing connections, leading to improvement in behaviour.

## INTRODUCTION

Cortical damage that accompanies traumatic brain injury or stroke often extends beyond the boundaries of the initial injury. This can lead to maladaptive cortical reorganisation and cognitive impairment (Grefkes & Fink, 2014). On the other hand, beneficial cortical reorganisation following injury can also occur and this can lead to recovery of function. Understanding the nature of cortical reorganisation after injury and how this might be promoted is a challenge for research on developing treatments for patients suffering from brain injury.

Resting state functional connectivity (rsFMRI) provides an indirect method of measuring cortical organisation across the whole brain by correlating BOLD activation patterns between pairs of brain areas. Strong correlation implies, at minimum, a “functional” connection, and often an anatomical connection (Deco, Jirsa, & McIntosh, 2011). Over the past decade, rsFMRI has been used to examine changes in network organisation in healthy individuals as well as patients who have suffered lesions or who have a variety of disorders such as schizophrenia, Alzheimer’s and Parkinson’s disease (Fornito, Zalesky, & Breakspear, 2015; He et al., 2007; Siegel et al., 2016). However, to fully understand the consequences of cortical reorganisation following damage, it is necessary to measure correlations within cortical networks both pre-and post-injury. This is rarely possible with human patients, and so we must rely on animal models, where we can collect data both before and after a lesion.

Recent studies have looked at functional connectivity following lesions in non-human primates. O’Reilly and colleagues (2013) sectioned the corpus callosum (with/without anterior commissure section) in monkeys to explore the relationship between structural connectivity and functional connectivity in neocortical areas. Grayson and colleagues (2016) used designer receptors exclusively activated by designer drugs (a.k.a. DREADDS) to temporarily inactivate the amygdala and were able to show how acute changes in functional connectivity in amygdala-cortical and cortico-cortical networks followed structural connectivity patterns. However, neither study related these changes to behaviour. By contrast, Meng and colleagues (2016) made neurotoxic lesions in the hippocampi of infant monkeys and correlated the resulting long-term changes in functional connectivity with performance on memory tests when the animals were 8-10 years old.

These studies are important in demonstrating the utility of studying functional connectivity following lesions. However, to understand how behavioural recovery occurs after brain injury in patients, we need to correlate changes in functional connectivity with behavioural measures as recovery occurs. In the present study, we therefore used rsFMRI to study how cortico-cortical connectivity in the macaque monkey brain changed in response to discrete lesions to regions near the principal sulcus (specifically, areas 46 and 9/46) of the prefrontal cortex over an extended period of time; and how these changes related to behaviour.

We chose to study lesions to regions near the principal sulcus because it is well known that lesions there reliably abolish the ability of monkeys to perform delayed response and delayed alternation tasks (E. K. Miller & Cohen, 2001; Passingham & Wise, 2012). This area has extensive anatomical connections to other frontal regions as well as with parietal and temporal regions (Petrides & Pandya, 1984; Saleem, Miller, & Price, 2014; Yeterian, Pandya, Tomaiuolo, & Petrides, 2012). We trained two monkeys on a location-based and object-based delayed match-to-sample task. We collected rsFMRI data at periodic intervals during the pre-lesion period to coincide with behavioural testing sessions. The animals then first received a lesion to both banks of the principal sulcus, including areas 46 and 9/46. Following a post-operative recovery period, we resumed periodic testing and scanning sessions. Later, they received a second lesion to the same region in the opposite hemisphere and they were once again tested and scanned at regular intervals. Adding the second lesion allowed us to assess the contribution to recovery of the homotopic region in the undamaged hemisphere.

## RESULTS

We performed a longitudinal assessment of the effect of lesions to the principal sulcus (areas 46 and 9/46) on behavioural performance on two cognitive tasks and related it to changes in functional connectivity (Figure 1). Once the animals had reached a predefined level of performance on the behavioural tasks (>70%), we collected functional neuroimaging (rsFMRI) data under general anaesthesia at two intervals prior to the first lesion, separated by 3-4 weeks (Data from two additional scans, earlier in the animals’ training, are not included in the present report). Several days prior to each scanning session, the animals were tested on both the location-and object-based delayed match-to-sample (DMS) tasks. Following these two cycles of behavioural testing and scanning, each animal received a lesion to both the dorsal and ventral banks of the left principal sulcus (PS), targeting areas 9/46, 46d, and 46v (Figure 2). Following a post-operative recovery period (approximately 4 weeks), we resumed cycles of behavioural testing and scanning (4 cycles approximately once/3-4 weeks). For the next several months, similar cycles of behavioural testing (but without scanning) continued. After 7 months following the first lesion, the animals received a second lesion to both banks of the right PS (Figure 2). Following a post-operative recovery period (approximately 4 weeks), the animals were once again tested and scanned (4 cycles, approximately once every 3-4 weeks).

**Figure 1:**
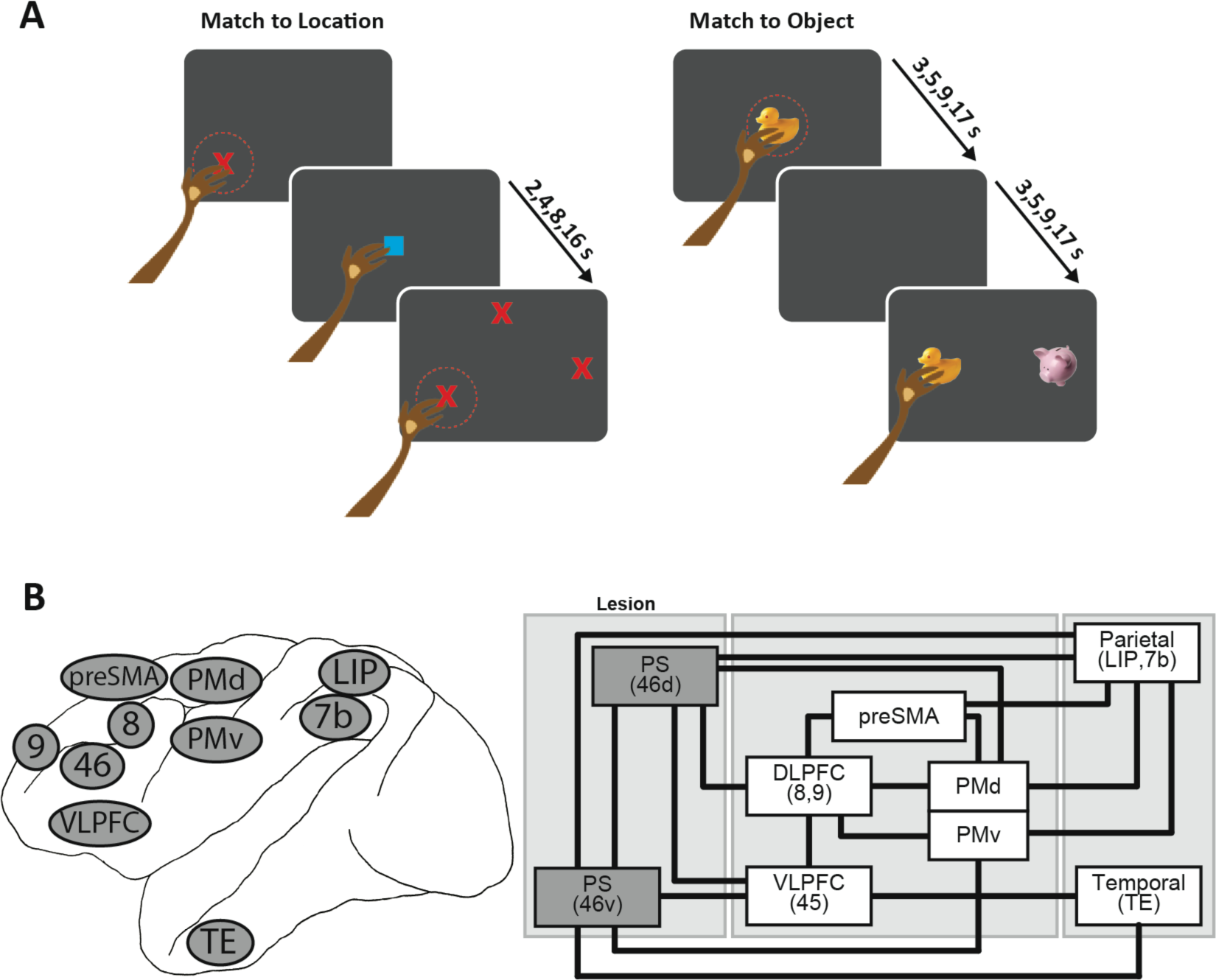
Assessing behavioural impairments and cortical reorganisation following lesions to the principal sulcus. A) Behavioural testing took place with the monkeys unrestrained in transport boxes, facing a touchscreen. In the ‘match-to-location’ task (left), the monkey was required to touch a cue that appeared in a random location on the touchscreen. The cue then disappeared and after a variable delay, three stimuli identical to the cue appeared in three different locations. The three locations included the sample location from the current trial, the cued location from a previous trial, and a third random location. The monkey was required to touch the location of the cue on the current trial to receive a food pellet reward. In the ‘match-to-object’ task (right), the monkey was once again required to touch a cue that appeared in a random location on the touchscreen. After a variable delay, two different stimuli appeared in random locations: the sample stimulus and a distracter stimulus. The monkey was required to touch the sample stimulus to receive a food pellet reward. B) Location and anatomical connectivity for regions in our network of interest. TE: area TE; PMd: dorsal premotor area; PMv: ventral premotor area; LIP: lateral intraparietal area; 7b, 8, 9, 46: areas 7b, 8, 9, 46; VLPFC: ventrolateral prefrontal cortex; preSMA: pre-supplementary motor area; PS: principal sulcus.

**Figure 2:**
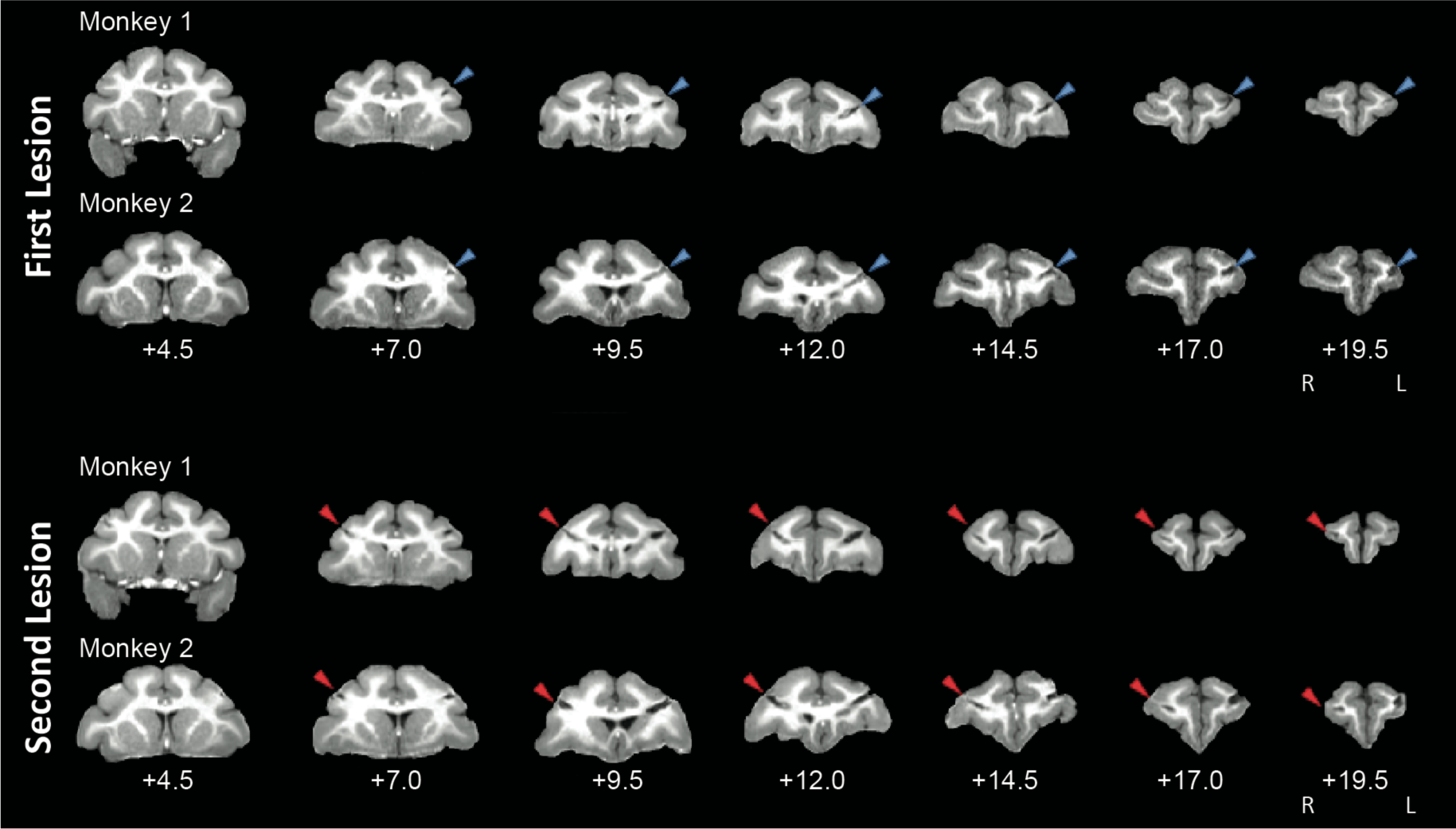
Structural MRIs showing location of first and second lesions in monkey 1 and 2. Coronal slices showing lesion to principal sulcus. Anatomical distances shown relative to the interaural axis.

### Behavioural deficits following unilateral lesion associated with decreased functional connectivity within frontal cortex

The ability of both monkeys to perform the two DMS tasks was significantly impaired following a unilateral lesion of the left PS regions (Figure 3A). We compared behavioural performance (measured as % correct) in the two sessions prior to the lesion (pre-lesion-1) with the first two sessions following the lesion (early post-lesion-1) and the two after 8 weeks (late post-lesion-1) using a mixed-model ANOVA (Figure 3A; see MATERIALS AND METHODS). We observed a significant main effect of monkey (*F*_(1,13)_=23.20, p=3.37×10^−4^), along with a significant interaction between monkey and experimental stage (*F*_(2,13)_=5.74, p=0.0164). We also observed a significant main effect of task (location vs. object, *F*_(1,13)_=87.34, p=3.92×10^−7^). Critically, a main effect of experimental stage (*F*_(1,13)_=17.90, p=1.84×10^−4^) was observed, including a notable decrease in performance shortly following the lesion, followed by substantial recovery. All remaining main effects and interactions failed to achieve statistical significance (p’s>0.05).

**Figure 3:**
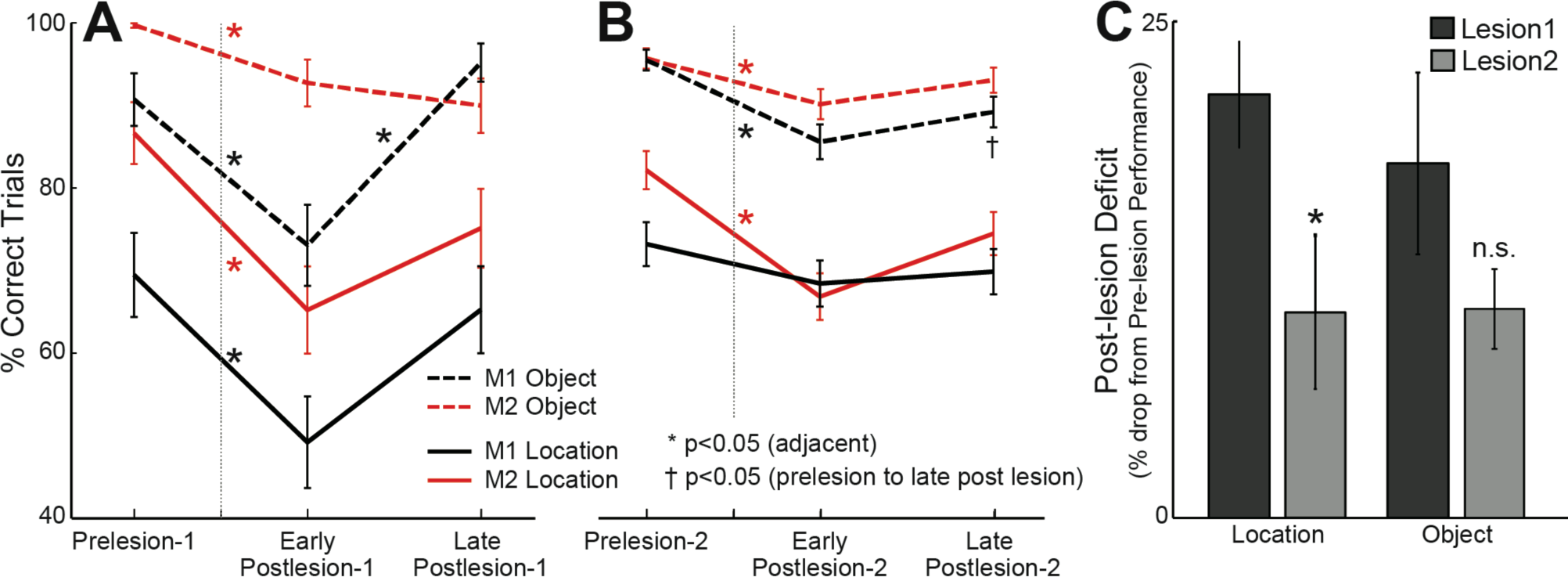
Behavioural performance of both monkeys in the match-to-location (solid lines) and match-to-object (dashed lines) tasks. Data are plotted as mean+/-SEM. A) Following a lesion to the left principal sulcus, both monkeys showed a significant drop in performance in both tasks in the testing sessions conducted within 8 weeks of the lesion. In the behavioural testing sessions beginning about 8 weeks post-lesion, both animals showed significant improvement. B) By contrast, after a second lesion to the right principal sulcus, performance in the location task was impaired in monkey 2 but not monkey 1. Performance in the object task for both monkeys decreased (*p<0.05, t-tests between adjacent time-points, ^†^p<0.05, t-tests between prelesion and postlesion-late timepoints; corrected for multiple comparisons). C) Impairments in the location and object tasks in the 8 weeks immediately following the first and second lesions (*p<0.05, paired t-tests).

Compared to their pre-lesion levels, both monkeys generated significantly fewer correct responses in both the location and object tasks in the early period following the lesion (Figure 3A). In the location task, the performance of monkey 1 decreased from 70±5% to 49±6% (mean±SEM, p=0.016); and the performance of monkey 2 decreased from 87±4% to 65±5% (p=0.0041). In the object task, the performance of monkey 1 decreased from 91±3% to 73±5% (p=0.0072); and the performance of monkey 2 decreased from 100±1% to 93±3% (p=0.0026, Holm-Bonferroni corrected t-tests).

This decrease in behavioural performance during the first 8 weeks following the lesion (early post-lesion-1) was associated with significant changes in functional connectivity within our network of interest. In Figure 4, we compare the changes in functional connectivity in the post-lesion periods to the pre-lesion levels, with the data grouped according to lobe and hemisphere (see Supplementary Figure 2 for changes to individual connections following the first lesion). We analysed these data using a mixed-model ANOVA (see MATERIALS AND METHODS). We observed significant main effects for experimental stage (*F*_(2,6)_=6.46, p=0.036) and connection (*F*_(20,60)_=28.62, p=1.96×10^−6^), and a significant interaction between the two (*F*_(4,24)_=4.954, p=0.00463). We did not observe a main effect of monkey (*F*_(1,6)_=1.27, p=0.302).

**Figure 4:**
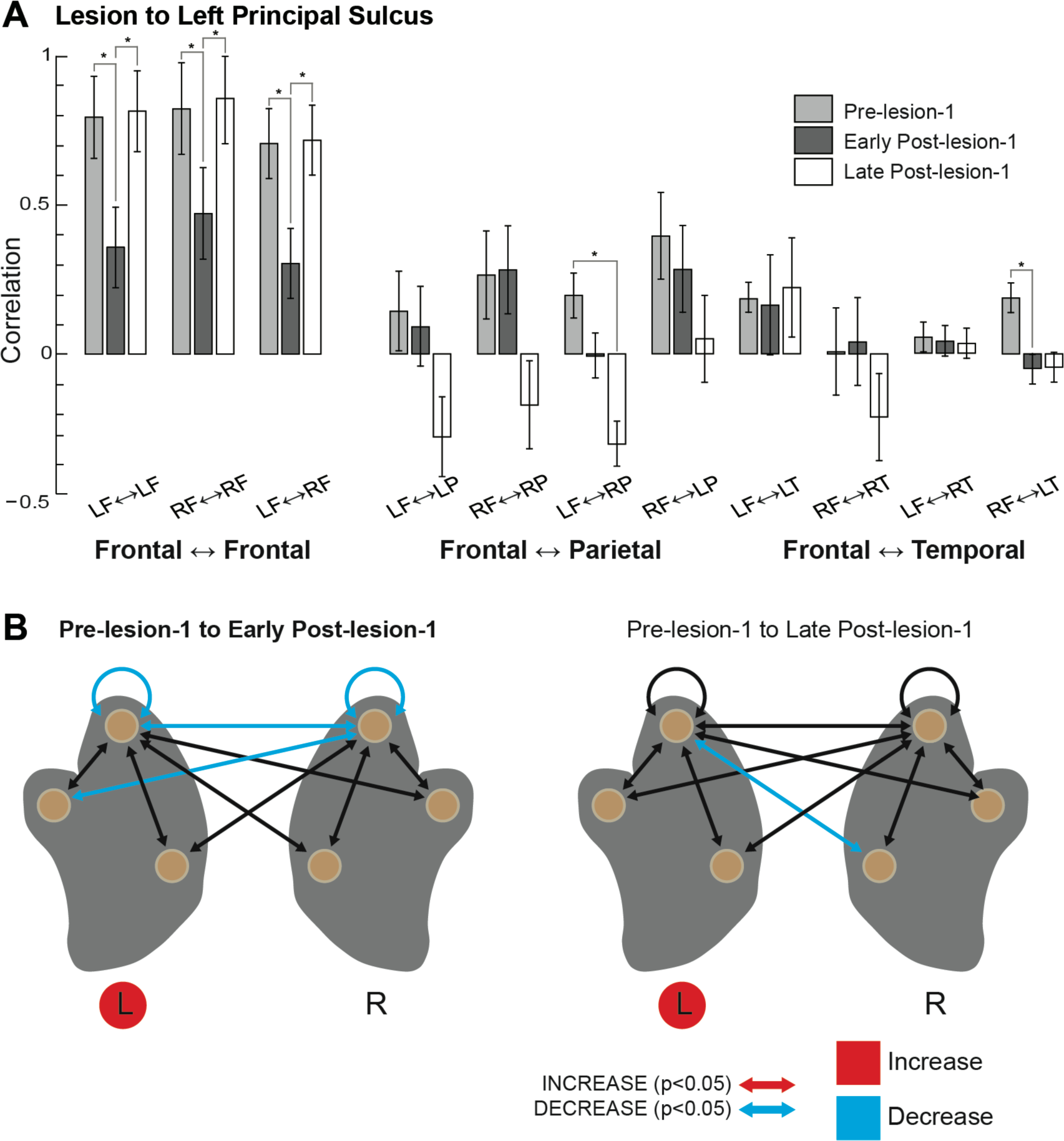
Connectivity within frontal lobes impaired following lesion to the left principal sulcus. A) Average correlation for all frontal-frontal, frontal-parietal, and frontal-temporal connections in the pre-lesion, early-post-lesion (4-8 weeks post lesion) and late post-lesion (8-16 weeks) periods. B) Schematic representations of changes to connectivity in the early post-lesion (left) and late post-lesion (right) periods, relative to the pre-lesion period. Significant increases in functional connection strength (correlation) shown in red; decreases in blue.

The most striking change in the first series of scans following the first lesion to the left PS was an overall decrease in correlation strength both within and between regions in the left and right frontal lobes (Figure 4). The correlations between frontal and parietal areas were unchanged (p’s>0.05). The correlations between frontal and temporal areas were also unchanged, except for the average connection between right frontal and temporal lobes (p=0.031; all the other frontal-temporal correlations p’s>0.05).

### Behavioural recovery associated with restored frontal connectivity

In later behavioural sessions conducted 8-12 weeks after the lesion (late postlesion-1), both monkeys showed signs of recovery on the location task, and in the case of monkey 1 on the object task (Figure 3A). On the location task, the performance of monkey 1 increased from 49±6% to 65±5% (p=0.069), and the performance of monkey 2 increased from 65±5% to 75±5% (p=0.28). For both monkeys, performance improved to the point where it was no longer statistically different from their pre-lesion performance (pre-lesion-1 vs. late postlesion-1, p=0.43 and p=0.095 for monkeys 1 and 2, respectively). On the object task, the performance of monkey 1 increased from 73±5% to 95±2% (p=0.006). It too was restored to pre-lesion levels (p=0.38). The performance of monkey 2 showed a marginal decrease from 93±3% to 90±3% (p=0.78).

At the same time, functional connectivity within the network of interest showed evidence of returning to a pre-lesion state. In Figure 4, we compare the changes in functional connectivity in our network of interest in the late post-lesion-1 period to the pre-lesion-1 period. At this stage, functional connectivity within and between frontal lobes had returned largely to the pre-lesion state.

### Disruption to behavioural performance following subsequent lesion to right principal sulcus

Approximately seven months following the lesion to the left PS regions, both monkeys received a second lesion to the right PS regions (Figure 2). We used a second mixed-model ANOVA to evaluate the behavioural impact of this procedure across the three stages of behavioural testing: the 8-week period immediately before the second lesion (pre-lesion), the first 8 weeks following the second lesion (early post-lesion-2), and the period 8-12 weeks following the second lesion (late post-lesion-2).

The behavioural data following the second lesion are shown in Figure 3B. We observed a significant main effect of monkey (*F*_(2,13)_=6.89, p=0.021), experimental stage (*F*_(1,13)_=17.06, p=2.24×10^−4^), and task (location vs. object, *F*_(1,13)_=207.99, p=2.24×10^−9^). All the interactions, including monkey and experimental stage, failed to reach statistical significance (p’s>0.05).

In general, removal of the previously-intact right PS regions had a smaller impact on behavioural performance as compared to the first lesion. On the location task, the performance of monkey 1 decreased from 73±3% in the pre-lesion-2 period to 68±3% in the early post-lesion-2 period (p=0.41); while the performance of monkey 2 decreased from 82±2% to 66±3% (p=0.0006). On the object task, the performance of monkey 1 decreased from 95±1% to 85±2% (p=0.0006), and the performance of monkey 2 decreased from 96±1% to 90±2%, (p=0.027).

We were unable to collect imaging data immediately prior to the second lesion. Therefore, to assess changes in connectivity associated with the second lesion, we compared functional connectivity from the scans performed after the second lesion to the latest scans after the first lesion. Thus, what we have called late post-lesion-1 is identical to what we now refer to as pre-lesion-2.

In Figure 5, we compare broad-scale changes in functional connectivity following the second lesion in the post-lesion periods to the pre-lesion levels, with the data grouped according to lobe and hemisphere (see Supplementary Figure 3 for changes to individual connections following the second lesion). As before, we analysed these data using a mixed-model ANOVA (see MATERIALS AND METHODS). Unlike the first lesion, the main effect for experimental stage failed to achieve statistical significance (*F*_(2,6)_=2.154, p=0.20). We did, however, observe a significant main effect for connection (*F*_(10,60)_=20.072, p=4.21×10^−7^) and a significant interaction between experimental stage and connection (*F*_(20,60)_=2.7798, p=0.028). We did not observe a main effect of monkey (*F*_(1,6)_=2.353, p=0.18).

**Figure 5:**
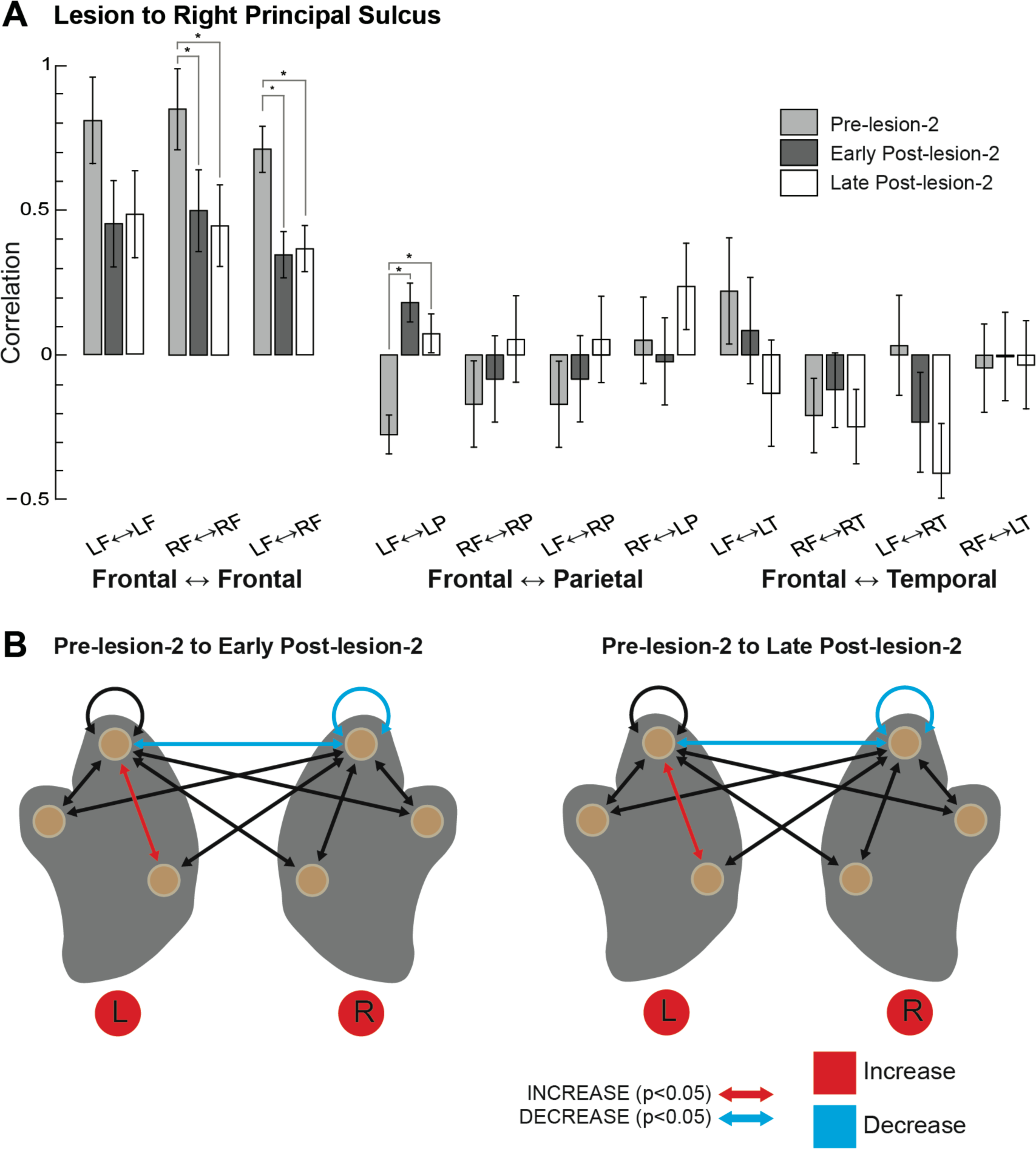
Connectivity within frontal lobes impaired following lesion to the right principal sulcus. A) Average correlation for all frontal-frontal, frontal-parietal, and frontal-temporal connections in the pre-lesion, early-post-lesion (4-8 weeks post lesion) and late post-lesion (8-16 weeks) periods. B) Schematic representations of changes to connectivity in the early post-lesion (left) and late post-lesion (right) periods, relative to the pre-lesion period. Significant increases in functional connection strength (correlation) shown in red; decreases in blue.

As with the lesion to the left PS regions, removing the right PS regions led to decreased correlations within and between the frontal lobes in the early post-lesion-2 as compared to the pre-lesion-2 periods. This was significant for the right hemisphere (pre-lesion-2 vs. early post-lesion-2, 0.80 ±0.15 vs. 0.37±0.15, p=0.019), and for the inter-hemispheric correlations (0.72±0.08 vs. 0.35±0.08, p=0.005). However, it did not reach statistical significance for correlations within the left hemisphere (0.74±0.15 vs. 0.34±0.15, p=0.0603). We observed a significant increase in average correlations between the left frontal and parietal lobes (pre-lesion-2 vs. early post-lesion-2, 0.27 ±0.07 vs. 0.18±0.07, p=0.019).

### Absence of network reorganisation or behavioural recovery in later post-lesion period

Behavioural performance did not change significantly from the early post-lesion-2 period for either monkey in the location task (Figure 5A; early post-lesion-2 vs. late post-lesion-2: monkey 1: 68±3% vs. 69±3%; monkey 2 67±3% vs. 74±3%, p=0.61 and p=0.10, respectively) or the object task (monkey 1: 86±3% vs. 89±2%; monkey 2: 90±3 vs. 93±2, p=0.47 and p=0.31, respectively).

Compared to the first lesion, the most striking feature of network reorganisation following the second lesion was the absence of recovery in the late post-lesion period (compare Figure 5 vs. Figure 4). Correlations showed few differences in the late postlesion-2 period as compared to the early post-lesion-2 period: the average right hemisphere and inter-hemispheric correlations remained significantly reduced compared to the pre-lesion levels and the correlations between left frontal and left parietal lobes remained increased relative to pre-lesion levels (pre-lesion-2 vs. late post-lesion-2, −0.28±0.07 vs. 0.07±0.07, p=0.017).

## DISCUSSION

We have examined changes in functional connectivity between regions in frontal, temporal, and parietal cortex following unilateral and bilateral frontal lesions, and compared this with changes in behaviour. Following a unilateral lesion to the left PS regions, both monkeys showed a significant impairment in performance on both the location-based and object-based DMS tasks. This was associated with a generalised decrease in connectivity between frontal regions, both within and between hemispheres. After about 8 weeks, both monkeys showed an improvement in behavioural performance that was associated with largely restored connectivity within frontal regions. Following a second lesion to the opposite hemisphere, both monkeys once again showed reduced connectivity within frontal-frontal connections, with a marginal effect on behavioural performance. Importantly, following the second lesion, there was no sign of recovery in either behaviour or network integrity.

On the one hand, the network effects of a focal frontal lesion were surprisingly widespread. Following a unilateral lesion, disruption extended not just to connections between undamaged frontal regions within the lesioned hemisphere, but to connections of these to the opposite frontal lobe, and even to connections exclusively within the undamaged hemisphere. On the other hand, disrupted connectivity was far from universal. In both hemispheres, connections of frontal to parietal and frontal to temporal cortex were largely unchanged, following either unilateral or bilateral frontal lesion. These results suggest that, following focal frontal lobe lesions, behavioural impairments and recovery could be specifically related to a widespread disruption of connectivity; restricted to the frontal lobes but widespread within those lobes. Similarly, recovery of behaviour following a unilateral lesion was associated with bilateral recovery of frontal connectivity.

In the following sections, we first acknowledge potential limitations in our approach. We then discuss why a unilateral lesion to PS regions might lead to disruptions in behaviour. Next, we consider how recovery of connections within and between the frontal lobes could help compensate for the effect of the lesion and thus lead to an improvement in behavioural performance. Finally, we discuss the marginal behavioural impairment and lack of network recovery following a second lesion to the opposite hemisphere.

### Limitations and issues of interpretation

We acknowledge several potential limitations to this study. First, we only tested two animals and no control animals were included. When designing studies in non-human primates, one must balance the need for adequate sample size with the ethical, logistical, and financial costs associated with the work. We did not anticipate a need for control animals because both animals were trained to the same criterion prior to the first lesion. In the absence of any lesion, there was no reason to suspect that the animals would exhibit significant changes in behaviour.

Second, we cannot be sure that underlying white matter was not affected during the surgical procedures even though an operating microscope was used. And indeed, there is evidence from a few sections that the lesion may have extended beyond the fundus of the sulcus (Fig. 2). However, the purpose of our study was not to determine the function of the cortical tissue near the principal sulcus, but rather to study functional recovery; and in stroke patients, white matter is always affected. Regardless of the extent to which underlying white matter was affected, we were nonetheless able to observe both behavioural impairment and recovery that correlated with changes in functional connectivity.

Third, after the second lesion there were changes in functional connectivity; yet there was only a marginal change in behaviour. One possible confound is a practice effect, in that as the experiment progressed the monkeys received more and more practice on the two tasks. However, it is important to note that both monkeys were trained to criterion prior to the first lesion and they were tested relatively infrequently following the lesion. It is therefore unlikely that practice alone can account for the behavioural recovery following the first lesion and the marginal effect on behaviour of the second lesion. We instead argue that it is more likely the monkeys adopted a different strategy for completing the task that allowed them to bypass the effects of the lesions (see below). Consistent with (but not definitively in support of) this suggestion is the observed increase in connectivity between the parietal and prefrontal cortex in the hemisphere (left) in which we placed the first lesion (Fig. 5).

Fourth, we collapsed the behavioural data across testing cycles and difficulty (i.e., delays); which raises some issue with interpretation. We chose to collapse these data in order to increase statistical power and to better align the behavioural data with the imaging data. It is possible that doing so obscured a potentially interesting effect of difficulty on functional connectivity. Unfortunately, this would be impossible to assess with the current dataset as the imaging data were collected with the animals under anaesthesia (meaning we could not tease out any effect of performance on functional connectivity). Nonetheless, for completeness, we show the complete behavioural dataset in supplementary material.

Finally, we acknowledge a potential issue of voxelsize. The problem is that neighbouring voxels may be supplied by the same vessels. This means that there may be an artefactual correlation between adjacent regions. To best avoid this problem, we chose as seed areas regions that were less likely to be supplied in this way (Fig. 1).

### Why is behaviour impaired following the first lesion?

The first lesion included areas 46v (ventral bank of the PS) and 46d (dorsal bank). Despite their anatomical proximity, these two regions have quite different sets of connections, particularly with areas outside of the frontal cortex (Petrides & Pandya, 1984; Saleem et al., 2014; Yeterian et al., 2012) (Figure 1B). Area 46d has connections to frontal areas 8 and 9; strong connections to parietal area 7, cingulate areas 23, 24 and 31; and relatively strong connections to the superior temporal gyrus (STG) and the dorsal banks of superior temporal sulcus (STS). Area 46v has connections to the ventrolateral prefrontal cortex (areas 12 and 45); and strong connections to the dorsal insula, the ventral bank of the STS, and the parietal area 7b.

We speculate that there are two networks that are relevant for performance on our tasks. The first is a ‘dorsal network’, comprised of the DLPFC including areas 8, 9, and 46d, and this has extensive connections with parietal cortex and is involved in spatial encoding. The second is a ‘ventral network’, comprised of the VLPFC including areas 12, 45, 46v, and 47, and this has extensive connections with the temporal cortex and is involved in object processing (Passingham & Wise, 2012). Thus, the PS lies at the anatomical intersection between these two functional networks, and is in an ideal position to integrate information about location and object identity. That it does indeed do so is indicated by single-unit recording studies that have shown that neurons within the principal sulcus can have both ‘what’ and ‘where’ properties (Hoshi, Shima, & Tanji, 1998; Rao, Rainer, & Miller, 1997).

When the tissue surrounding the PS is damaged, we should therefore expect to see impairment in both spatially-based and object-based tasks, and this is indeed what was observed here and is consistent with the results of previous studies (Funahashi, Bruce, & Goldman-Rakic, 1993; Soper, 1979). Our results further suggest, however, that behavioural impairments should not be explained by the damage to area 46 alone. Instead, this damage broadly disrupts frontal lobe function, reflected in widespread reductions in frontal connectivity. Such widespread disruptions may arise precisely because area 46 itself is so broadly connected to other frontal regions, including both dorsal and ventral networks.

### How did recovery of function occur following the first lesion? Why was there only a marginal deficit despite a persistent disruption to network connectivity following the second lesion?

Both monkeys eventually showed recovery of function after the first lesion, and this brings us to the fundamental issue, which is how does this recovery occur. First, we must consider the possibility that the lesions were not complete and it was residual tissue that was mediating behavioural recovery. However, as can be seen from Figure 2, there was very little, if any, tissue left in the principal sulcus. It is one advantage of working with animals that it is possible to ensure, as here, that the lesions are indeed complete.

A second possibility is that after the first unilateral lesion, the homotopic region in the other hemisphere could compensate. But if area 46 in the right hemisphere had indeed taken over, we might have expected to see a severe impairment when it was then itself lesioned, which was not the case (Figure 3C); so at best this may only be part of the answer. A third possibility, and one that is most strongly supported by the data, is that widespread cortical recovery and reorganisation within the frontal lobes contributed to the recovery of function following the first lesion. Given the complexity of the cortical network and the degree of interconnectivity, particularly within and between the frontal lobes, there are always potential alternative routes of information transmission so long as the lesion is not extensive. If there are multiple potential strategies to complete the task, for example, it is conceivable that alternative pathways that bypass area 46 might be recruited, or might recover, as frontal connectivity is restored.

Results following the second lesion were somewhat mixed. The behavioural impairment was modest in comparison to the large impairment that followed the first, unilateral lesion and yet, there was little evidence of subsequent recovery. The disruption of frontal lobe connectivity was again substantial and this time long-lasting or permanent. Potentially, behaviour is only mildly affected by a second lesion to the right area 46 because by this point, the animals have either adopted a strategy that is less dependent on area 46 and/or alternative neural pathways that bypass area 46 have been recruited. Moreover, assuming this is true, there would be little drive for cortical reorganisation in this case, which would account for why little change in network connectivity was observed even at the late post-lesion-2 period. A second possibility is that, for frontal networks to recover, area 46 is necessary in at least one hemisphere, relating to the common clinical observation that especially severe cognitive deficits can follow bilateral frontal lesions. In this case, modest behavioural impairment following the second lesion might be ascribed to some additional factor, such as the additional training received between the two lesions.

### Summary and future applications

In line with many previous suggestions, our data confirm that the effects of a focal frontal lobe lesion cannot be understood simply as loss of function in the specific area removed. Instead, there is a widespread, though also specific, disruption of connectivity between many regions within and between the two frontal lobes, potentially bringing a widespread impairment in their function. At least following unilateral lesion, this disruption recovers over time, with associated recovery in behaviour. We suggest that it is only possible to understand how a brain lesion affects behaviour by studying the whole network and its interconnections. A lesion, however discrete, disturbs the network as a whole and this effect can last for weeks. However, recovery can occur as intact parts of the network regain their normal function, providing alternative ways in which the system can perform the task. Given the complexity of the network, there are multiple ways in which information in one area can be transmitted to inform another.

With this study, we have provided a framework for investigating recovery after lesions that relies on a non-invasive methodology (rsFMRI) that could be used with human patients. Critically, this methodology allows for longitudinal tracking of cortical reorganisation. Given enough data, this methodology could be used to generate predictions about clinical outcomes based on markers of cortical reorganisation following stroke, as in the study by (Siegel et al., 2016). The use of an animal model such as ours could be used to develop new therapies for promoting recovery in patients by driving cortical reorganisation. For example, cortico-cortical paired associative stimulation (ccPAS) has been used to enhance functional connectivity between visual areas leading to increased visual perception (Romei, Chiappini, Hibbard, & Avenanti, 2016) and motor plasticity (Chao et al., 2015; Johnen et al., 2015). However, fully evaluating the potential of techniques such as paired stimulation in driving *targeted* post-lesion recovery requires assessment of baseline connectivity prior to and post lesion. Such an assessment is feasible, as in our study, in a non-human primate model.

## MATERIALS AND METHODS

All animal procedures were conducted in accordance with the United Kingdom Animals (Scientific Procedures) Act (1986) and approved by the University of Oxford local ethical review panel and the UK Home Office Animal Inspectorate. All husbandry and welfare conditions complied with the guidelines of the European Directive (2010/63/EU) for the care and use of laboratory animals. Two adult male monkeys (*Macaca mulatta*, 8–11 kg), purpose-bred in the United Kingdom, were used in this study. The monkeys were pair-housed with varying forms of environmental enrichment, free access to water, and a 12-hour light/dark cycle. Veterinary staff performed regular health and welfare assessments, which included formalized behavioural monitoring.

### Behavioural tasks

Behavioural testing took place with the monkeys unrestrained inside small transport boxes (approximately 1 m^3^) (see (A. S. Mitchell, Baxter, & Gaffan, 2007) for details). One side of the testing box faced a touchscreen to which the monkey had access. In the ‘match-to-location’ task (Figure 1A, left), the monkey was required to touch a red cross that appeared in a random location on the touchscreen. The cross then disappeared and a distractor (blue square) appeared in the centre of the screen and the monkey was required to touch this. After a variable delay (2, 4, 8, or 16 s), three stimuli identical to the sample appeared in three different locations. The three locations included the sample location from the current trial, the sample location from a previous trial, and a third random location. The monkey was required to touch the cued location on the current trial to receive a food pellet reward.

In the ‘match-to-object’ task (Figure 1A, right), the monkey was required to touch a cue that appeared in the centre of the touchscreen. There was then a variable delay (3, 5, 9, or 17 s); the extra 1 second was added to approximately match the distractor plus delay durations in the match-to-location task. Two stimuli then appeared on the touchscreen on either side of midline (along the horizontal meridian, equidistant from centre). These included the sample stimulus and a distracter stimulus (randomly allocated to either left or right of midline). The monkey was required to touch the stimulus that had been cued to receive a food pellet reward.

The two tasks were not matched for overall difficulty: based on performance data, the location task was more difficult than the object task (Figure 3). For each testing cycle, there were 1 or 2 test sessions per task (100-120 trials per session), on different days. The second test session was added from the second postlesion cycle onwards. For cycles with 2 test sessions per task, data from the two were combined as there was no significant difference in performance between the sessions when summed across all tasks, monkeys, and stages of testing (testing session 1 vs. testing session 2: 82±2% vs. 83±2, p=0.11, paired t-test).

Because of the relatively long delays between testing sessions (several weeks), we started each cycle with shorter ‘warm-up’ sessions (40-100 trials). These were held over two days prior to actual testing sessions in order to re-introduce the animals to the process of testing. Data obtained during warm-up days were excluded from all analyses.

All performance data (percent correct) were arcsine transformed before being analysed using two separate 3-way mixed-model ANOVAs (one per lesion) with each testing cycle corresponding to a unit of replication. Each ANOVA included three fixed-effects: task (match-to-location, match-to-object), subject (monkey 1, monkey 2), and experimental stage (pre-lesion-1/2, early postlesion-1/2, late postlesion-1/2); and one random-effect corresponding to testing cycle. Post-hoc t-tests were carried out on ANOVA-derived estimated marginal means to identify specific changes in performance associated with subject, task and experimental stage. All p-values were adjusted for multiple comparisons using the Holm-Bonferroni method to control for family-wise error (Holm, 1979).

For presentation of data and reporting of summary statistics, arcsine-transformed values were back-transformed to yield values corresponding to percent correct.

### Neuroimaging Data Collection

All imaging data were collected under general anaesthesia on a 3T scanner using a custom-made 4-channel phased array coil (H. Kolster, MRI Coil Laboratory, Laboratory voor Neuro en Psychofysiologie, KU Leuven). Procedures for inducing and maintaining general anaesthesia and the positioning of the monkeys in the scanner are similar to those described previously (Mars et al., 2011; D. J. Mitchell et al., 2016).

Resting-state echo-planar images were collected at 2x2x2mm resolution (36 axial slices, TR=2 s, TE=19 ms). We collected between 825 and 1600 volumes per scan session (mean: 1511, SD: 223), for an approximate scan duration of 56 min. During this time, anaesthesia levels (isofluorane, ~0.8-2.0%, mean 1.53%) and respiration rate were held constant.

T1-weighted, high-resolution (0.5 mm isotropic voxels) structural images were collected using an MPRAGE (magnetization prepared gradient echo; TR = 2.5s, TE = 4.01 ms, 3-5 averages) sequence. Anatomical images were corrected for coil inhomogeneity by dividing the MPRAGE data by a lower resolution MPRAGE sequence (1x1x1 mm) that did not include an inversion recovery pulse (Parameters: TR = 2.5 s, TE = 3.48 ms) (Van de Moortele et al., 2009).

### Neurosurgery

Lesions to the principal sulcus (areas 46 and 9/46) were performed under aseptic conditions using an operating microscope (for a detailed description of the surgical procedures, see (Buckley & Mitchell, 2016; Buckley et al., 2009). To protect against intraoperative oedema and postoperative inflammation, steroids (methylprednisolone, 20 mg ⁄ kg, im) were administered the night before and three additional times 4–6 h apart (iv or im) on the day of surgery. Anaesthesia was initiated with a single dose of ketamine (10 mg/kg) and xylazine (0.25–0.5 mg/kg, im) and maintained using sevoflurane (1.0–2.0% to effect, in 100% oxygen) while the animal was mechanically ventilated. The animal was given atropine (0.05 mg/kg) to reduce secretions, an antibiotic (amoxicillin, 8.75 mg/kg) as prophylaxis against infection, and buprenorphine (0.01 mg/kg iv, repeated two times at 4-to 6-h intervals on the day of surgery iv or im) and meloxicam (0.2 mg/kg iv) for analgesia. Ranitidine, an H_2_ receptor antagonist (1 mg/kg iv), was given to protect against gastric ulceration as a side effect of the combination of steroid and nonsteroidal anti-inflammatory treatment. Heart rate, oxygen saturation, mean arterial blood pressure, end tidal CO_2_, body temperature, and respiration rate were monitored continuously throughout the procedure.

Under deep anaesthesia, the head was placed in a head holder and the skull exposed by opening the scalp and galea in layers. The temporal muscles were retracted and a bone flap was removed. The dura was cut to expose the cortical surface. Both banks of the principal sulcus were removed with aspiration (Figure 2). The dura was then sewn and the bone flap replaced.

Following the procedure, the animals were monitored continuously for 48 hours. Postoperative medication continued in consultation with veterinary staff. This included nonsteroidal anti-inflammatory analgesic (meloxicam, 0.2 mg ⁄ kg, oral) and antibiotic (8.75 mg ⁄ kg, oral) treatment following surgery in consultation with veterinary staff, typically for 5 days. In addition, steroids (dexamethasone, 1 mg/kg, im), once every 12 h for 3 days, then once every 24 h for 2 days; analgesia (buprenorphine, 0.01 mg/kg, im) for 48 h; and continued antibiotic treatment (amoxicillin, 8.75 mg/kg, oral) were also administered for 5 days. Gastric ulcer protection (omeprazole, 5 mg/kg, oral and antepsin, 500 mg/kg, oral) commenced 2 days prior to surgery and continued postoperatively for the duration of other prescribed medications, up to 5 days.

### Data Preprocessing and Analysis

All data preprocessing and analysis was conducted using a combination of Matlab (The MathWorks Inc.), SPM8 (Statistical Parametric Mapping; www.fil.ion.ucl.ac.uk/spm), FSL (fMRI of the Brain (FMRIB) Software Library; <u>http://fsl.fmrib.ox.ac.uk/fsl/fslwiki/;</u> (Jenkinson, Beckmann, Behrens, Woolrich, & Smith, 2012), Caret (Computerized Anatomical Reconstruction Toolkit; (Van Essen, 2012) and aa software (automatic analysis; (Cusack et al., 2014); <u>www.automaticanalysis.org</u>). The methods used to analyse rsFMRI data are described in detail in (D. J. Mitchell et al., 2016). Briefly, the structural volumes for each animal were aligned to standard space (112 Rhesus macaque template – in the space of the atlas of (Saleem & Logothetis, 2006) and segmented into grey matter, white matter and cerebrospinal fluid (CSF) tissue classes (Mclaren et al., 2009). After removing the first 6 functional volumes, the remaining volumes from each rsFMRI dataset were used to estimate motion parameters (included as covariates of no-interest) and were aligned to standard space through a two-stage process. The data were spatially-smoothed with a 3 mm Gaussian kernel (full-width half maximum). Grey-matter masks were defined as voxels with grey-matter probability >0.5 within each subject. Masks representing white matter, CSF, and the superior sagittal sinus were also produced and used to create covariates of no interest in time series analysis (see following).

Physiological covariates of no interest were constructed from the EPI time-series for white-matter and CSF tissue masks (up to 6 principal components each) (Behzadi, Restom, Liau, & Liu, 2007). A further vascular component was defined as the mean time-course within a mask drawn for the superior sagittal sinus. The first temporal derivatives of these time-courses were also included. A motion confound covariate was defined as the time-course of average displacement over the expected brain volume (approximated as a sphere of radius 40 mm). The first temporal derivative of this vector was also included, as were the element-wise squares of both vectors. Discrete cosine transform covariates were added to implement a temporal band-pass filter (0.0025-0.05Hz) as commonly used (Mantini et al., 2011; Vincent et al., 2007). After projecting covariates from each grey-matter time-series, functional connectivity (“connection strength”) was estimated as correlations between the mean grey-matter time-series, for a range of pre-defined regions of interest (ROIs). ROIs were derived from the macaque cortical parcellation scheme proposed by Van Essen and colleagues (Van Essen, Glasser, Dierker, & Harwell, 2012).

### Defining the network of interest

To evaluate changes in connectivity following the lesions, we compared pre-and post-lesion connectivity in a predefined “network of interest” (shown in Figure 1B) composed of a subset of frontal, parietal, and temporal regions. These regions were selected on the basis of their anatomical connectivity to areas 46 and 9/46 (Saleem et al., 2014; Yeterian et al., 2012), specifically focusing on those that have been implicated in the performance of location-and object-based cognitive tasks (Passingham & Wise, 2012). These regions were delineated according the macaque cortical parcellation scheme proposed by Van Essen and colleagues (Van Essen et al., 2012). They included in the frontal cortex: dorsal area 6DR (“PMd”), ventral area 6Val (“PMv”), medial area 6M (“preSMA”), areas 8Ac and 9 (areas 8 and 9, collectively referred to as “DLPFC”), area 45 (“VLPFC”). They included in the parietal cortex: area 7b, dorso-and ventro-lateral intraparietal areas (LIPd, and LIPv). Finally, they included in the temporal cortex: TE1-3d (“TE”). A schematic of the intrahemispheric anatomical connectivity between these regions and areas 46 and9/46 is shown in Figure 1B (based on (Saleem et al., 2014; Yeterian et al., 2012).

### Analysing changes in functional connectivity

To analyse broad-scale changes in connectivity, we averaged the different pairwise connections according to hemisphere and lobe. This yielded 11 different values per subject and stage of the experiment, corresponding to all possible pairwise combinations of hemisphere and lobe [e.g., left-frontal to left-frontal (LF-LF), left-frontal to right-frontal (LF-RF), left-frontal to left-parietal (LF-LP), etc.]. Note that LF-RF and RF-LF yield identical values and so only a single group is used to summarise these data. The data were modelled as a fixed effect in two separate mixed-model ANOVAs (one per lesion). These ANOVAs also included two further fixed-effect factors [experimental stage (3 levels, pre-lesion-1/2, early postlesion-1/2, late postlesion-1/2); monkey (2 levels)] and 1 random-effect (replication sessions within each experimental stage). Post-hoc t-tests were carried out on ANOVA-derived estimated marginal means (corrected for multiple comparisons using the Holm-Bonferroni method).

## ACKNOWLEDGEMENTS / FUNDING

This work was supported by the MRC intramural program MC-A060-5PQ10 (MA, DM, JD, AB), an MRC Career Development Award G0800329 (AM). The authors thank Jerome Sallet, Matthew F Rushworth, Richard N Henson for advice on connectivity analysis; Subhojit Chakraborty with data collection; Stuart Mason for training the animals, and the Oxford Biomedical Sciences Staff for their assistance with the animals. The authors declare no competing financial interests.

---

**Supplementary Figure 1.**
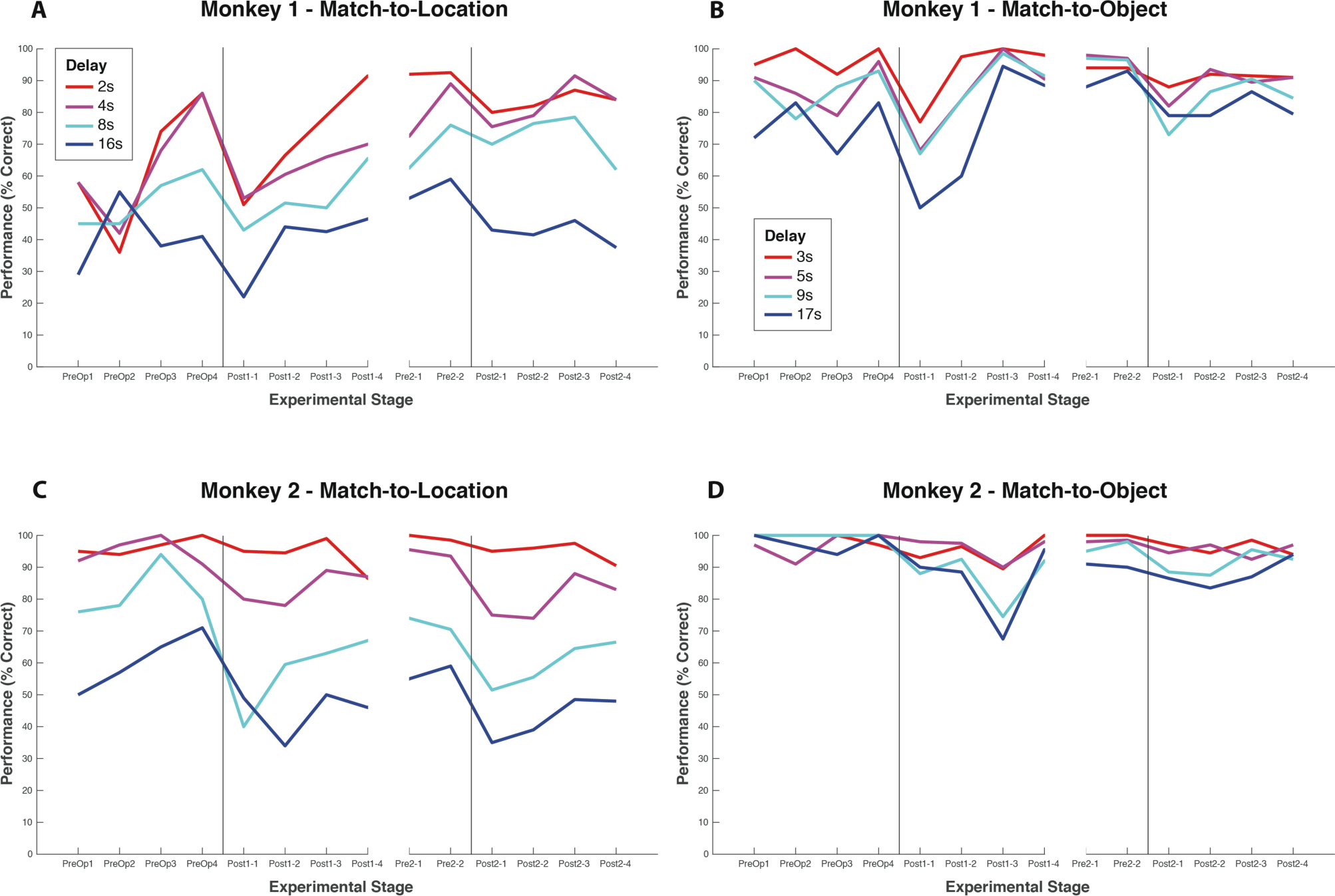
Behavioural performance of both monkeys in the match-to-location (A,C) and match-to-object (B,D) tasks. Data are plotted as percent correct within individual testing sessions, separated according to delay between cue and probe stimuli.

**Supplementary Figure 2.**
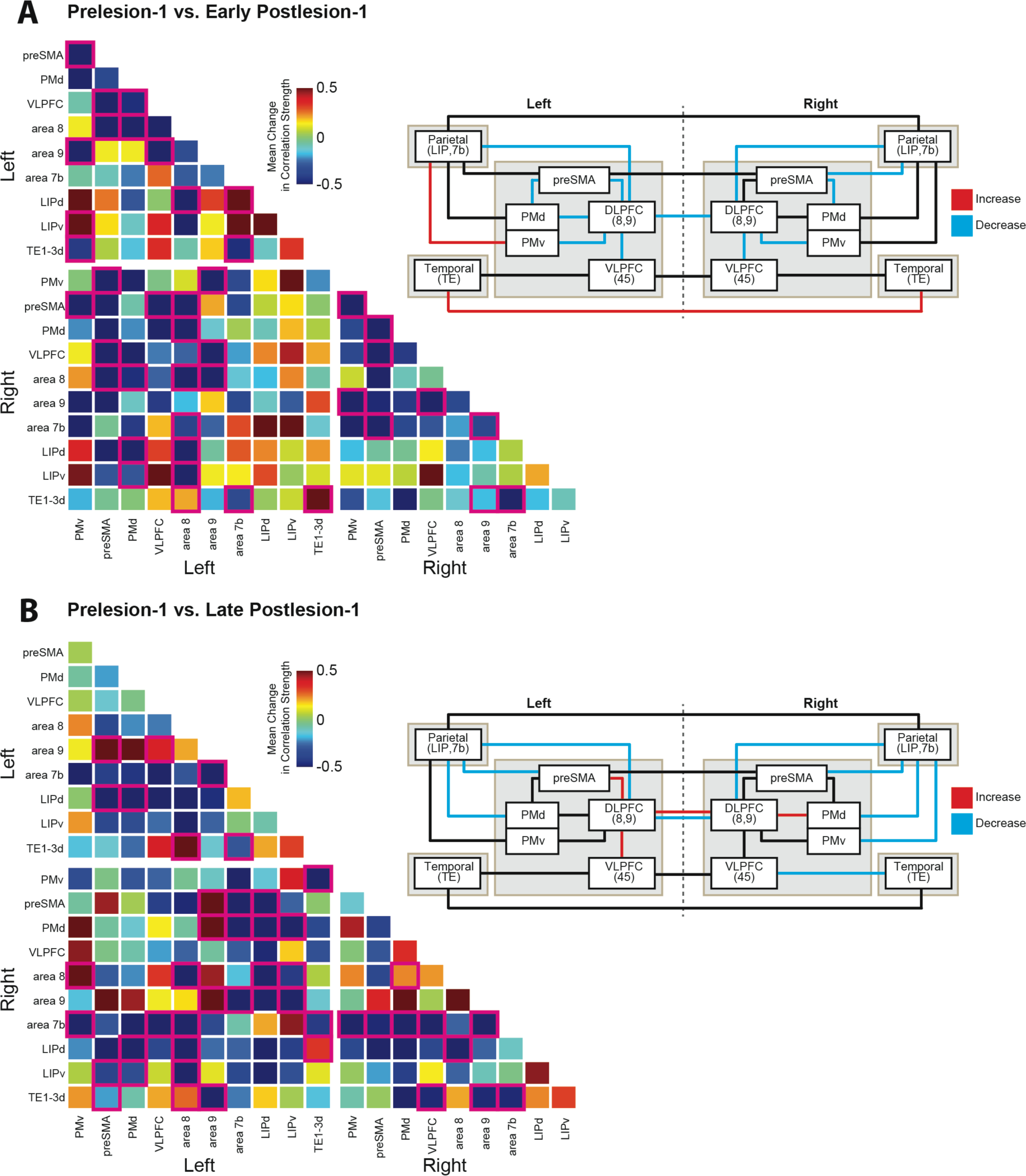
Changes in pairwise connectivity within network of interest following lesion to left areas 46 and 9/46. Connectivity matrices showing changes in functional connectivity (as measured by correlation strength) between pairs of regions within the network of interest for the early post-lesion (A) and late post-lesion (B) periods, as compared to the pre-lesion dataset. Squares outlined in pink indicate those pairwise connections that fell outside a 99% confidence interval of connection strength across that pairwise comparison in the pre-lesion dataset. To the right of each matrix is a schematic representation of the changes to connectivity compared to the pre-lesion dataset. Increases in functional connection strength shown in red; decreases in blue. Connections are coloured if any of the possible connections between the nodes show a change in connection strength outside the 99% confidence interval.

**Supplementary Figure 3.**
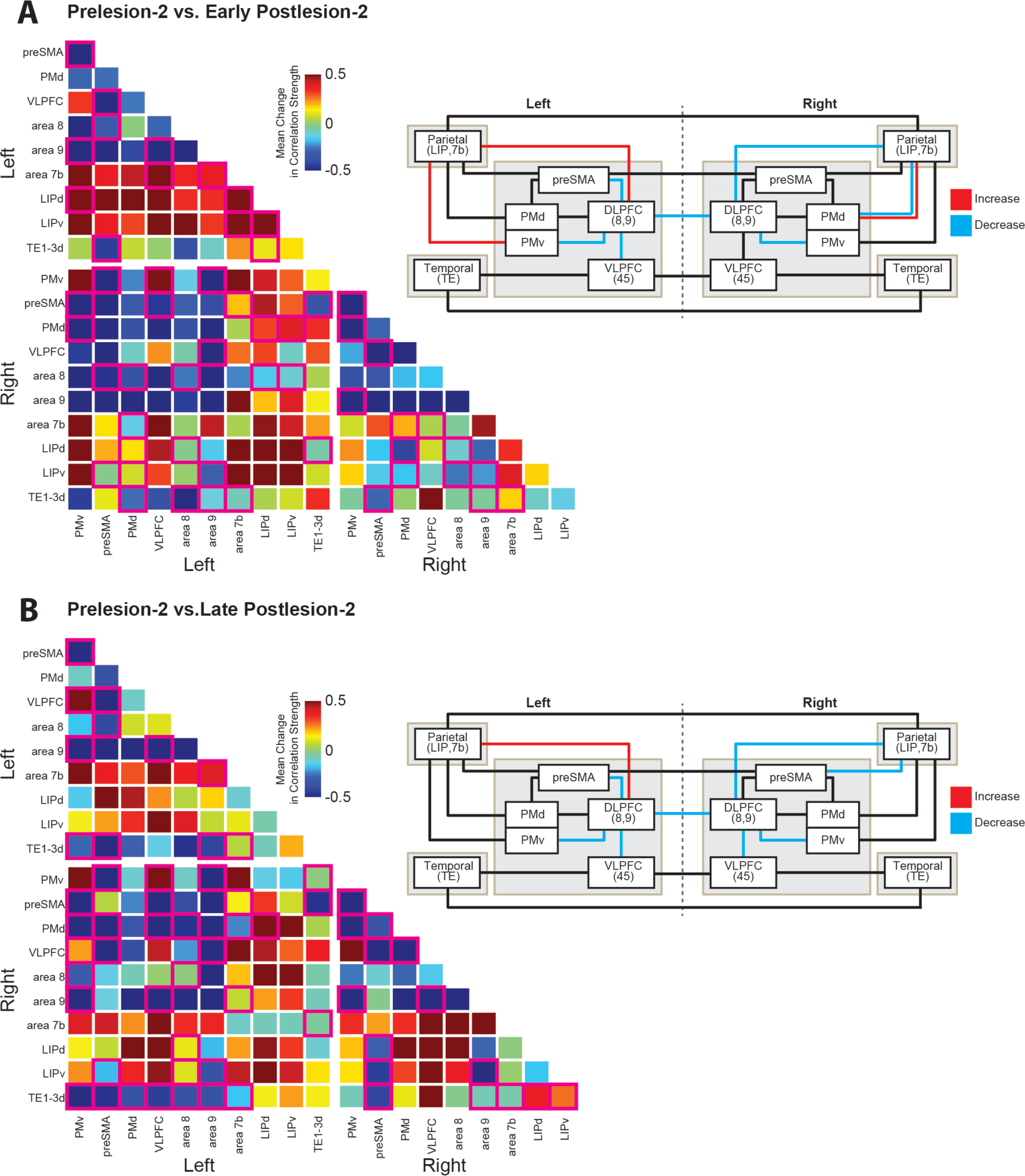
Changes in pairwise connectivity within network of interest following lesion to right areas 46 and 9/46. Connectivity matrices showing changes in functional connectivity (as measured by correlation strength) between pairs of regions within the network of interest for the early post-lesion (A) and late post-lesion (B) periods, as compared to the pre-lesion dataset. Squares outlined in pink indicate those pairwise connections that fell outside the 99% confidence interval of connection strength across that pairwise comparison in the pre-lesion dataset. To the right of each matrix is a schematic representation of the changes to connectivity compared to the pre-lesion dataset. Increases in functional connection strength shown in red; decreases in blue. Connections are coloured if any of the possible connections between the nodes show a change in connection strength outside the 99% confidence interval.

